# Vaping and smoking cue reactivity in young adult electronic cigarette users who have never smoked combustible cigarettes: A functional neuroimaging study

**DOI:** 10.1101/2024.01.13.575524

**Authors:** Jiaying Liu, Zhenhao Shi, Jessica L. Fabbricatore, Joshua T. McMains, Allison Worsdale, Erin C. Jones, Yidi Wang, Lawrence H. Sweet

## Abstract

**Introduction:** The rapid growth in the use of electronic cigarettes (e-cigarettes) among young adults who have never smoked combustible cigarettes is concerning, as it raises the potential for chronic vaping and nicotine addiction. A key characteristic of drug addiction is the elevated neural response to conditioned drug-related cues (i.e., cue reactivity). Generalized reactivity to both vaping and smoking cues may signify an increased risk for smoking initiation in non-smoking vapers. In this study, we used functional magnetic resonance imaging (fMRI) to evaluate brain responses to vaping and smoking cues in young adult never-smoking vapers.

**Methods:** Sixty-six young adult never-smoking vapers underwent functional MRI while viewing visual cues pertaining to vaping, smoking, and nicotine-unrelated unconditioned reward (i.e., food). A priori region-of-interest analysis combined with exploratory whole-brain analysis was performed to characterize neural reactivity to vaping and smoking cues in comparison to food cues.

**Results:** The medial prefrontal cortex and the posterior cingulate cortex, regions that play a key role in drug cue reactivity, showed significantly increased neural response to vaping cues compared to food cues. The posterior cingulate cortex additionally showed increased neural responses to smoking cues compared to food cues.

**Conclusions:** Despite never having smoked combustible cigarettes, young adult vapers exhibited heightened neural susceptibility to both vaping and smoking cues within brain systems associated with cue reactivity. The findings shed light on the mechanisms underlying nicotine addiction and smoking initiation risk in this critical population and may contribute to the development of science-based interventions and regulatory measures in the future.

**IMPLICATIONS:** The escalating vaping prevalence among US never-smoking young adults is alarming, due to its potential ramifications for nicotine addiction development. Nicotine addiction is characterized by elevated neural response to conditioned nicotine-related cues. Using functional neuroimaging, we showed that young adult non-smoking vapers exhibited heightened neural susceptibility to both vaping and smoking cues within brain systems previously associated with cue reactivity. Such cross-reactivity to both types of nicotine cues may serve as the mechanism underlying nicotine addiction and smoking initiation risk in this population. Our findings may contribute to the development of science-based interventions and regulatory measures addressing the vaping epidemic.

## INTRODUCTION

The use of electronic cigarettes (e-cigarettes) is rapidly growing among adolescents and young adults. Despite its potential to facilitate the cessation of combustible cigarette smoking ^1^, vaping is paradoxically most prevalent among young adults who do not use e-cigarettes as a means to quit smoking ^2,3^. Chronic vaping leads to nicotine addiction, which increases the risk for vaping escalation and smoking initiation ^4,5^.

A key characteristic of drug addiction is elevated psychophysiological responses to conditioned drug-related stimuli, i.e., cue reactivity ^6^. After repeatedly signaling the delivery of drug rewards, stimuli that are otherwise neutral to drug-naïve individuals may acquire high incentive value and trigger drug seeking among users. Functional magnetic resonance imaging (fMRI) studies show that drug cue reactivity is mediated by a number of cortical and subcortical brain regions that underlie the processing of motivational salience, including the medial prefrontal cortex (MPFC), anterior and posterior cingulate cortex (ACC, PCC), anterior insula (AI), nucleus accumbens (NAcc), caudate, and amygdala ^7,8^. Among combustible cigarette smokers, heightened neural response to smoking cues in these systems has been linked to future smoking behavior and cessation outcomes ^9,10^. Emerging evidence also shows increased brain reactivity to vaping cues among e-cigarette users ^11,12^.

Given that smoking and vaping cues both signal nicotine availability, vapers with established nicotine addiction may exhibit cross reactivity to both types of nicotine cues even if they are not yet accustomed to combustible cigarette smoking. Heightened response to smoking cues in the brain’s motivational salience circuitry may reflect an increased likelihood of smoking initiation for never-smoking vapers. The aim of the current study was to examine the neural mechanisms of vaping and smoking cross cue reactivity using fMRI. We hypothesized that the brain cue-reactivity circuits will show greater response to vaping and smoking cues compared to natural reward (food) cues in young adult vapers who never smoke.

## METHODS

### Participants

Sixty-six never-smoking young adult vapers participated in the study (see **Table 1** for participant characteristics). Potential participants were screened for study eligibility and fMRI compatibility over the phone. Participants were required to be between 18 and 29 years of age, have vaped 15 or more of the past 30 days, and have never smoked cigarettes. Exclusion criteria included history of major medical, psychiatric, or neurologic conditions and research contraindications for fMRI, including, but not limited to, some metal implants, pregnancy, or claustrophobia.

**Table 1.** Participant characteristics.

| Variable | mean±SD or count |
| --- | --- |
| Age (years) | 20.08±1.50 |
| Sex | 20 male, 46 female |
| Race | 49 European American, 3 African American,<br>9 Asian, 5 multi-race/other |
| Hispanic ethnicity | 8 Hispanic, 58 non-Hispanic |
| Nicotine dependence for e-cigarettes<sup>1</sup> | 14 no dependence, 18 low dependence,<br>19 medium dependence, 15 high dependence |
| No. of vaping days during past 30 days | 24.71±7.41 |
| <sup>1</sup> Dependence was measured by the Penn State Electronic Cigarette Dependence Index. |  |

Participants gave written informed consent to participate in the protocol. The study was approved and monitored by the Institutional Review Board of the University of Georgia.

### Procedure

Participants first completed a survey that included measures of sociodemographic variables and the Penn State Electronic Cigarette Dependence Index (PS-ECDI) ^13^. The PS-ECDI is a 10-item questionnaire that measures vaping-specific nicotine dependence levels. After the survey, participants completed a one-hour fMRI scan session that included a cue reactivity paradigm. Specifically, participants underwent two 6 min 54s runs of the fMRI visual cue-reactivity task. During each run, they viewed three 30-second blocks of images representing each of the three conditions: vaping, smoking, and food cues. Within each block, six distinct images were displayed for 5 seconds each. There were three 20-second blocks featuring non-rewarding control images (i.e., office products) and three 20-second blocks of scrambled images that served as baselines. The presentation order of the blocks was pseudorandomized. Participants were asked to rate their craving for e-cigarettes, cigarettes, and food, respectively, on a 7-point Likert scale (1=Not at all, 7=Very much) on four occasions during each run.

### MRI data acquisition and preprocessing

MRI data were collected at the University of Georgia using a GE Discovery 750 3T MRI scanner and a 32-channel head coil (GE Healthcare, Milwaukee, WI). Blood oxygenation level-dependent (BOLD) fMRI was performed, using the whole-brain, single-shot gradient-echo echo-planar pulse sequence with the following parameters: repetition time (TR)/echo time (TE)=2000/25 ms, field of view (FOV)=225×225 mm^2^, matrix=64×64, slice thickness=3.5 mm, flip angle (FA)=90°, 40 axial slices, effective voxel resolution of 3.52×3.52×3.50 mm^3^. After BOLD fMRI, structural images were acquired using the 3D fast spoiled gradient-echo sequence with the following parameters: TR/TE=7.80/2.98 ms, FOV=256×256 mm^2^, matrix=256×256,164 sagittal slices, FA=20°, effective voxel resolution of 1×1×1 mm^3^.

### MRI data analysis

MRI data were preprocessed and analyzed using SPM 12 (Wellcome Trust Centre for Neuroimaging, London, UK) and ArtRepair (www.nitrc.org/projects/art_repair) toolboxes in MATLAB (MathWorks, Natick, MA). MRI images underwent slice-wise outlier detection and repair (using art_slice), slice time correction, realignment and head motion correction, coregistration of structural and functional images, normalization into the Montreal Neurological Institute (MNI) space with 3-mm cubic voxels via unified segmentation, spatial smoothing by an 8-mm full-width at half-maximum Gaussian filter, and volume-wise outlier correction (using art_global). Trials were modeled by the canonical hemodynamic response function and its time derivative in a general linear model, which generated session-specific parameter estimate maps for each stimulus category and the rating periods. Six rigid-body motion parameters were included as covariates. Neural response to vaping, smoking, and food cues were evaluated by linear contrasts against baseline.

Brain regions that have been consistently activated by nicotine-related cues were defined as *a priori* regions of interest (ROIs) using the Neuromorphometrics atlas (neuromorphometrics.com): MPFC, ACC, PCC, AI, NAcc, caudate and amygdala ^7,8^ (see **Figure 1A**). Mean contrast values representing cue categories versus the scrambled control baseline were extracted from each ROI and subjected to analysis of variance (ANOVA) that examined the differences in neural response across cue categories. Because seven ROIs were tested, p-values of the ANOVAs were corrected for false discovery rate (FDR) following the Benjamini-Hochberg procedure.Significant ROIs were subjected to pairwise comparisons between cue categories using Tukey’s test.

**Figure 1.**
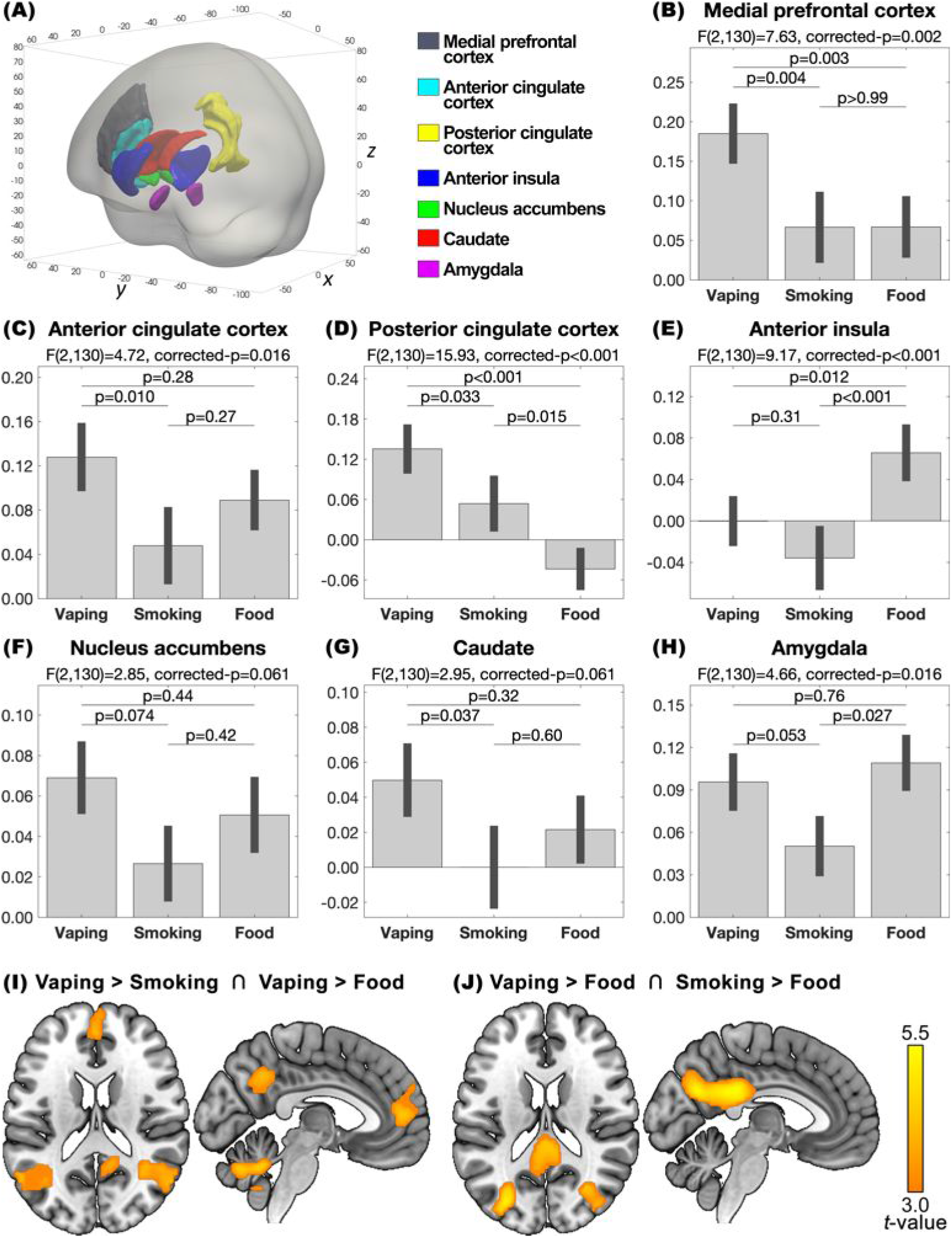
(Panel A) Definition of *a priori* regions of interest. (Panels B-H) Results of the region-of-interest analyses on the contrast values for the neural responses to vaping, smoking, and food cues compared to control stimuli. P- values for the F-tests were false discovery rate-corrected for the test of 7 regions. P-values for the pairwise comparisons were corrected using Tukey’s method. (Panel I) Results of the whole-brain conjunction analysis showing brain regions that had greater neural response to vaping cues than smoking and food cues. (Panel J) Result of the whole-brain conjunction analysis showing brain regions that had greater neural response to vaping and smoking cues than food cues. Significant clusters were identified at voxel-level p<0.001 and cluster-level false discovery rate-corrected p<0.05.

Group-level whole-brain ANOVAs were performed for each voxel on the contrast images to explore brain regions other than the ROIs that exhibited differential neural response to cues. Voxels that exhibited significant main effect of cues were used to generate an inclusive mask for the subsequent pairwise comparisons. To examine brain regions that showed greater reactivity specifically to vaping cues than both smoking and food cues, we performed a conjunction analysis for “vaping>smoking” ∩ “vaping>food” ^14^. To examine brain regions that showed greater reactivity to nicotine cues in general (i.e., vaping and smoking cues) than food cues, we performed a conjunction analysis for “vaping>food” ∩ “smoking>food”. Significant regions were determined using a voxel-level threshold of uncorrected p<0.001 combined with cluster-level FDR-corrected p<0.05.

## RESULTS

Participant characteristics are summarized in **Table 1**.

*A priori* ROI analyses showed a significant main effect of all cue types on the neural response in the MPFC, ACC, PCC, AI, and amygdala (partial η^2^=0.11, 0.07, 0.20, 0.12, & 0.07, FDR-corrected p’s<0.016), but not the NAcc or the caudate (partial η^2^=0.04 & 0.04, FDR-corrected p’s>0.061) (see **Figure 1B-H**). Post hoc comparisons revealed that both the MPFC and the PCC exhibited greater response to vaping cues than smoking and food cues, with the PCC additionally showing greater response to smoking than food cues. The AI showed greater response to food cues than vaping and smoking cues. There was also greater ACC response to vaping than smoking cues, and greater amygdala response to food than smoking cues.

Whole-brain ANOVA and conjunction analysis showed significantly greater reactivity specifically to vaping cues compared to smoking and food cues in the MPFC (cluster extent=5,562 mm^3^, Z=4.39, x/y/z=–3/50/8), PCC (cluster extent=8,370 mm^3^, Z=4.05, x/y/z=–3/–55/38), left inferior parietal lobule (cluster extent=7,884 mm^3^, Z=4.39, x/y/z=–60/–61/20), right inferior parietal lobule (cluster extent=13,770 mm^3^, Z=4.51, x/y/z=60/–61/8), and cerebellum (cluster extent=11,907 mm^3^, Z=4.37, x/y/z=–3/–55/–28) (see **Figure 1I**). We also found significantly greater reactivity to both types of nicotine cues (i.e., vaping and smoking) compared to food cues in a cluster that included the PCC (cluster extent=71,037 mm^3^, Z=5.29, x/y/z=–3/–40/26), medial superior parietal lobule (a.k.a. precuneus) (Z=5.21, x/y/z=–6/–64/26), left superior parietal lobule (Z=5.54, x/y/z=–27/–73/50), and right superior parietal lobule (Z=4.31, x/y/z=30/– 67/47) (see **Figure 1J**).

## DISCUSSION

We found that young adult never-smoking vapers had greater MPFC and PCC response to vaping-related visual cues compared to smoking and food cues. The MPFC and PCC are among the key brain regions underlying drug cue reactivity in individuals with substance use disorders ^9,10^. Among healthy individuals, the MPFC and PCC engage in the processing of information that is of high self-relevance ^15^ as well as envisioning one’s future behavior ^16^.These findings suggest increased incentive salience of vaping cues and high self-relevance of vaping behavior among vapers.

The PCC additionally showed significantly cross reactivity to smoking cues, albeit not as pronounced as its response to vaping cues. The finding is intriguing given that the participants only used e-cigarettes and never smoked cigarettes. Cross cue reactivity has previously been reported among established poly-substance users (e.g., alcohol + cannabis) ^17^. The observed smoking cue reactivity among non-smoking vapers may reflect generalized neurocognitive susceptibility to signals of nicotine availability, which are assigned a high incentive value as a result of vaping-induced nicotine addiction ^4^. Given the involvement of the PCC in self-referential and future-oriented thinking ^15,16^, PCC response to smoking cue reactivity may serve as a mechanism of piqued interest in experimenting smoking that could trigger smoking initiation among never-smoking vapers.

Exploratory whole-brain analysis showed that, in addition to the MPFC and PCC ROIs, the bilateral inferior parietal lobule and cerebellum also exhibited vaping-specific cue reactivity. Furthermore, regions adjacent to the PCC, including the medial and bilateral superior parietal lobule, showed cross reactivity to both vaping and smoking cues. The inferior parietal lobule is considered a part of the default mode network together with the MPFC and PCC and is involved in a wide range of functions including attention, memory, multisensory and linguistic processing, self-reflection, and social cognition ^18^. The medial and lateral superior parietal lobule are similarly multifunctional regions implied in diverse perceptual, cognitive, and motor processes ^19^. Future research is needed to clarify the exact role of the inferior and superior parietal lobe in nicotine cue reactivity.

Our study comes with several limitations. First, although our sample size was relatively large for neuroimaging ^20^, it may not be sufficient to detect small effect sizes. Notably, the striatal ROIs, i.e., the NAcc and caudate, showed a marginally significant main effect of cues that did not survive the FDR correction for multiple ROI comparisons. Increasing the sample size will help characterize the neural response patterns in the striatal cue-reactivity regions and allow for the testing of more subtle effects in additional brain areas. Second, the absence of control cohorts limited our ability to determine whether the results were specific to young adult non-smoking vapers. It will be interesting should future studies elucidate the cross-cue reactivity phenomenon in diverse populations that vary by age, vaping status, and smoking status. Lastly, the current study did not attempt to associate neural activity measures with behavioral measures, which prevented us from answering important questions regarding the behavioral and clinical implications of the fMRI findings. Some of the a priori ROIs exhibited cue reactivity patterns that were inconsistent with our hypotheses: the AI showed a selective response to food cues only, whereas the amygdala appeared to have shown a selective reduction in response to smoking cues. Incorporating longitudinal behavioral measures in future research may facilitate the interpretation of the unexpected neuroimaging discoveries. It will also help determine whether neural cue reactivity measures are predictive of future vaping and smoking behaviors.

In conclusion, our fMRI data showed that in young adult e-cigarette vapers who never smoked combustible cigarettes, exposure to vaping-related visual cues compared to natural reward cues increased the neural response of the MPFC and PCC, regions that are previously implicated in drug cue reactivity ^7,8^. The PCC additionally showed heightened response to smoking cues compared to natural reward cues. This is the first study to demonstrate increased neural susceptibility to both vaping and smoking cues in vapers who never smoke. The findings shed light on the neurocognitive mechanisms underlying the increased risk of smoking initiation in this critical population and may contribute to science-based regulation of the sales of e-cigarette products to non-smoking individuals.

## FUNDING

Data collected in this study was supported by the National Institutes of Health (NIH) grant K01DA049292, University of Georgia (UGA) Internal Junior Faculty Seed Grant in STEM, and UGA Owens Institute for Behavioral Research and Bioimaging Research Center Pilot Grant. JL was supported by the NIH (K01DA049292; R21DA056570). ZS was supported by the Brain & Behavior Research Foundation (NARSAD Young Investigator Grant #30780) and the NIH (K01DA051709). This research was also supported by an endowment from Gary R. Sperduto.

## DECLARATION OF INTERESTS

None declared.

## AUTHOR CONTRIBUTIONS

JL and LHS designed the study. JL, JLF, JTM, AW, ECJ, YW, and LHS collected the data. JL, ZS, and LHS performed the analysis and interpreted the results. JL, ZS, and LHS wrote the manuscript. All authors critically reviewed the manuscript and approved the final version for publication.

## DATA AVAILABILITY

The data that support the findings of this study are available from the corresponding authors upon reasonable request.

## ACKNOWLEDGEMENTS

The authors acknowledge the contribution of our undergraduate research assistant Nina Bhatikar, Emma Jones, Nidhi Manikkoth, Adhya Chawla, Neha Gregory, Saba Alemayehu, Ryan Quisling, Timothy Coy and Kiyan Kojoori for assisting in participant recruitment and data collection.

## Notes

### Competing Interest Statement

The authors have declared no competing interest.

